# *In situ* observation and range extension of the first discovered monoplacophoran *Neopilina galatheae*

**DOI:** 10.1101/2023.11.08.566290

**Authors:** Chong Chen

## Abstract

Monoplacophoran molluscs have been dubbed “living fossils” due to their absence in the fossil record for about 375 million years, until *Neopilina galatheae* Lemche, 1957 was trawled off Costa Rica in 1952. Since then, over 35 species of living monoplacophorans have been discovered. Nevertheless, *in situ* observations of these rare deep-sea animals remain scant. Here, we observed and collected an intact specimen of *N. galatheae* using a remotely operated vehicle from 2460 m deep on the Eastern Galápagos Spreading Center. The animal was found attached to the glassy surface of solidified basalt lava flow, and no feeding trails were found near the animal. Such hard substrate is in contrast with previous records that were trawled on sand and mud, suggesting *Neopilina* can be found on a wide range of substrates. This is the first time this species was collected since 1959, and represents a southeast range extension of about 1000 km for the species.

## Main Text

Monoplacophoran molluscs were thought to be extinct since the Devonian (∼375 million years ago) – until 1952 when suddenly 10 live individuals were brought up by the Danish *Galathea* expedition from 3570 m deep off Costa Rica in a single trawl on muddy clay (Lemche, 1957; Lemche & Wingstrand, 1959). Named *Neopilina galatheae* Lemche, 1957, the discovery of this first living monoplacophoran mollusc is widely regarded as one of the most important zoological discoveries of the 20^th^ Century (Lindberg, 2009; Ponder, Lindberg & Ponder, 2020). The Galathea trawl was a true “jackpot”, as follow-up cruises set to obtain further specimens near the type locality returned empty handed (Wolff, 1961) except the research vessel (R/V) Vema, which successfully collected a single specimen from 3718 m deep in 1958 (Menzies *et al*., 1959). Then, three live specimens turned up off Baja California, Mexico in 1959 where they were trawled between 2781–2809 m deep, plus a fragment from a grab sampler at 1828 m (Parker, 1961; Wolff, 1961). These were taken from sandy sediment rich in organic matter, foraminifera tests, and quartz (Parker, 1961). And that was the last definitive record of the mythical “living fossil” *N. galatheae*, with no new specimen being collected for over six decades (Schwabe, 2008).

Since the discovery of *N. galatheae*, over 30 living species of monoplacophorans have been described from around the globe (Ponder *et al*., 2020). Most of these are small species below about 5 mm shell length. At a maximum recorded length of 37 mm, *N. galatheae* remains the largest species (Lemche & Wingstrand, 1959). Some small species have been found as shallow as 177 m (Wilson *et al*., 2009), but large species in the genera *Neopilina, Vema*, and *Adenopilina* have only been found in waters over 1800 m deep (Parker, 1961; Tebble, 1967). Although some small monoplacophorans have been recovered alive and observed in aquaria (Lowenstam, 1978) there is only one convincing seafloor observation of a living monoplacophoran in its natural habitat, that of an undescribed *Neopilina* species found off American Samoa (Sigwart *et al*., 2019). These sightings were of animals living on basalt and associated with potential feeding trackways Some early seafloor images of the Atacama Trench from the *Vema* expedition contained straight trackways on mud that were claimed to be trails of the monoplacophoran *Vema ewingi* (Clarke & Menzies, 1959; Menzies *et al*., 1959). However, these trackways have been suggested to be more likely a misidentification of bivalve trails (Wolff, 1961) and their maker remains debated.

Here, we report a new record of *N. galatheae* in the Galápagos Spreading Centre, representing a range extension to the south and also providing a first glimpse into the natural habitat of this ‘living fossil’. Seafloor observation and sample collection were carried out using Schmidt Ocean Institute’s 4500 m rated remotely operated vehicle (ROV) SuBastian during cruise FKt231024 on-board R/V Falkor (too). Imagery was carried out using a 4K video camera (SULIS Subsea Z70; resolution 3840 × 2160 pixels) with 12X zoom capability. A CTD (conductivity, temperature, depth) sensor (Seabird FastCAT SBE49) and a dissolved oxygen Sensor (Aanderaa 3841 O2 Optode) on the ROV took real-time measurements at 1 second intervals. Sampling was done using a suction sampler mounted on ROV SuBastian.

Upon recovery of the ROV, the monoplacophoran specimen was retrieved from the suction chamber and gently cleaned using a brush. The cleaned specimen was observed with a Leica S APO stereomicroscope and photographed under chilled seawater using a EF 100 mm F2.8L MACRO IS USM macro lens and a Canon EOS 5Ds R digital single-lens reflex camera. Measurements were taken using a vernier calliper. Tissue snips were carefully taken from the posterior half of the foot using an iris scissor for future molecular work. After that, the entire monoplacophoran specimen was preserved in 80% ethanol. The specimen and all tissue snips are deposited in the Senckenberg Natural History Museum, Frankfurt under the catalogue number SMF 373198, with a mitochondrial cytochrome oxidase *c* subunit I (COI) barcode (Folmer *et al*., 1994) available on NCBI GenBank with the accession number PP352643.

Dive #603 of ROV SuBasitan was conducted at the Rose Garden hydrothermal vent field on the Galápagos Rift where deep-sea hydrothermal vents were first discovered in 1977 (Corliss *et al*., 1979). An eruption event between 2005 and 2012 is known to have covered this area with a vitreous fresh lava flow, leading to the cessation of almost all venting activity (Shank *et al*., 2012). During this dive ropey or lobate basaltic lava with glassy surface were seen across the area, often covered by a very thin layer of sediment (Figure 1A). There was a complete lack of ongoing hydrothermal venting and only a patch of empty, dissolving serpulid tubes and empty shells of mussels and clams indicated the presence of diffuse flow not so long ago.

**Figure 1.**
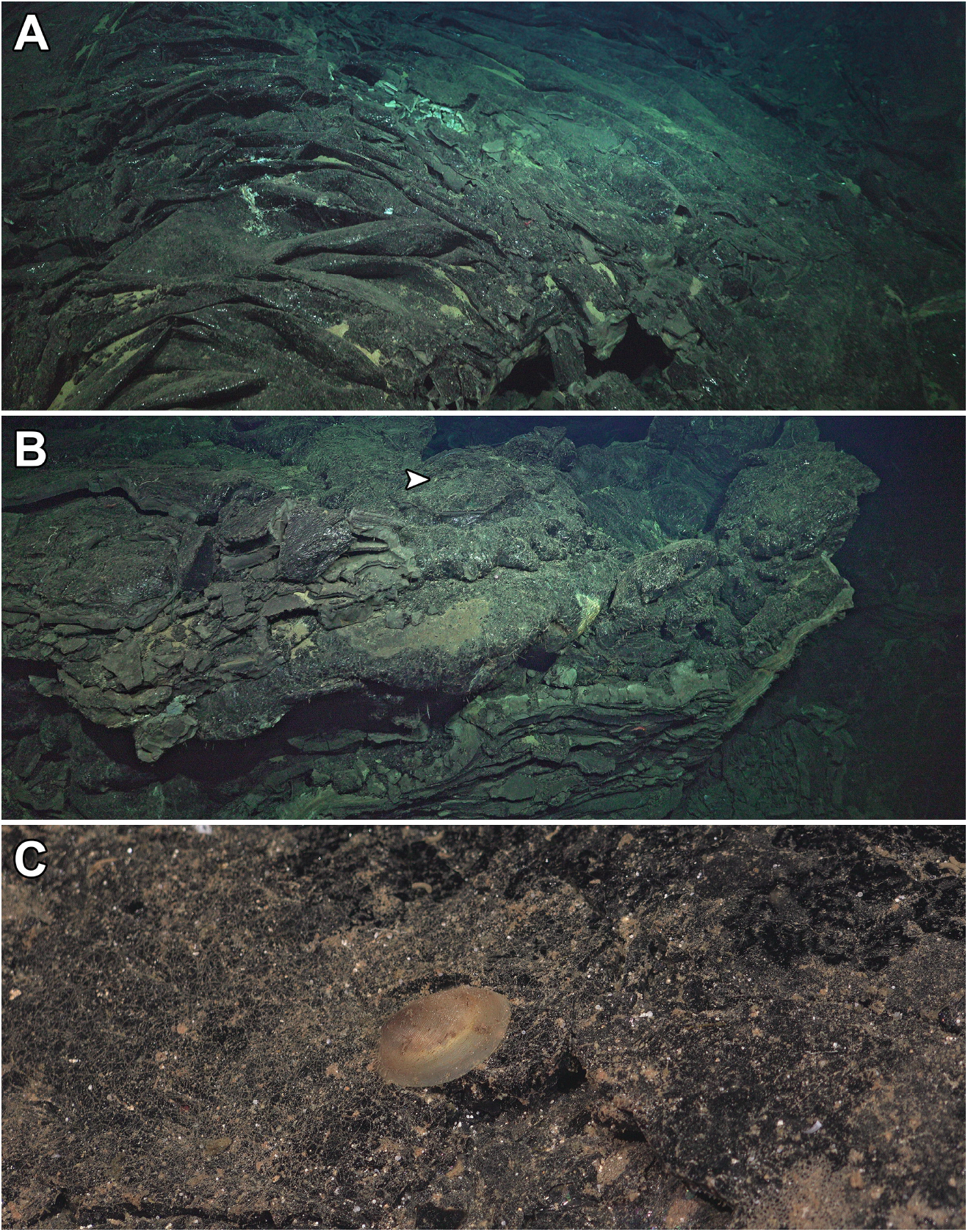
(A) Typical substrate seen in Rose Garden, the collecting site, consisting of rather fresh, vitreous lobate and ropey lava flows. (B) The lava tower from which the *Neopilina galatheae* specimen was collected, white arrow indicates the location of *N. galatheae*. (C) Close-up on the *N. galatheae* individual in life position.

Near the top of a half-collapsed lava tower (Figure 1B; 0°48.2986’N, 86°13.1732’W, 2461 m deep), we found a limpet-like animal (Figure 1C). Based on the very thin, translucent bilaterally symmetrical shell carrying a characteristic irregular or wavy growth lines, this was identified as a large-sized monoplacophoran (Lemche, 1957; Sigwart *et al*., 2019). The monoplacophoran individual lived attached to the surface of a glassy basalt flow covered by a thin layer of sediment littered with numerous foraminiferal tests and hydroids. No feeding trails could be observed around the animal, and it did not move during the course of our observation which lasted approximately five minutes. To collect this extremely fragile specimen, the pilot first used the ROV’s manipulator arm equipped with soft rubber on the tips to gently break the glassy lava around and below the monoplacophoran. After dislodging the specimen, it was then collected using suction sampler. A supplementary video available on Figshare (Chen, 2024) details the collecting process. The monoplacophoran was collected at 00:46 UTC on October 28th, 2023 (water temperature 2.09°C, salinity 34.656, dissolved oxygen concentration 3.337 mg/L). The area had no hydrothermal influences.

The monoplacophoran individual (Figure 2) was retrieved fully intact, but it was no longer responsive when recovered on surface. The shell length was 26.8 mm and the shell width was 24.0 mm. This large size, and the presence of five pairs of gills in the pallial groove at this size, points to genus *Neopilina* – as *Vema*, the only other genus known to reach this size, has six pairs of gills as adults (Clarke & Menzies, 1959; Lemche, 1957; Warén & Gofas, 1996). The two large-sized *Neopilina* species inhabiting the eastern Pacific, *N. galatheae* Lemche, 1957 and *N. bruuni* Menzies, 1968, are readily distinguished by the postoral tentacles which are significantly reduced in *N. bruuni* compared to *N. galatheae* (Menzies, 1968). The well-developed postoral tentacles in the newly collected specimen (Figure 2) and the matching gill structures as well as shell sculpture serve to identify this specimen as *N. galatheae*. It is medium-sized for *N. galatheae* (Lemche & Wingstrand, 1959), and as a result the apex position is not as anterior as the largest specimens; tracing growth lines in fig. 3 of Lemche & Wingstrand (1959) indicate that the holotype of *N. galatheae* would have had a similar apex position at a size comparable to this specimen. The gills, velum, postoral tentacles, outer edge of the foot, and pallial margin all carried bright orange pigmentation; while the lips around the mouth were reddish brown in colour.

**Figure 2.**
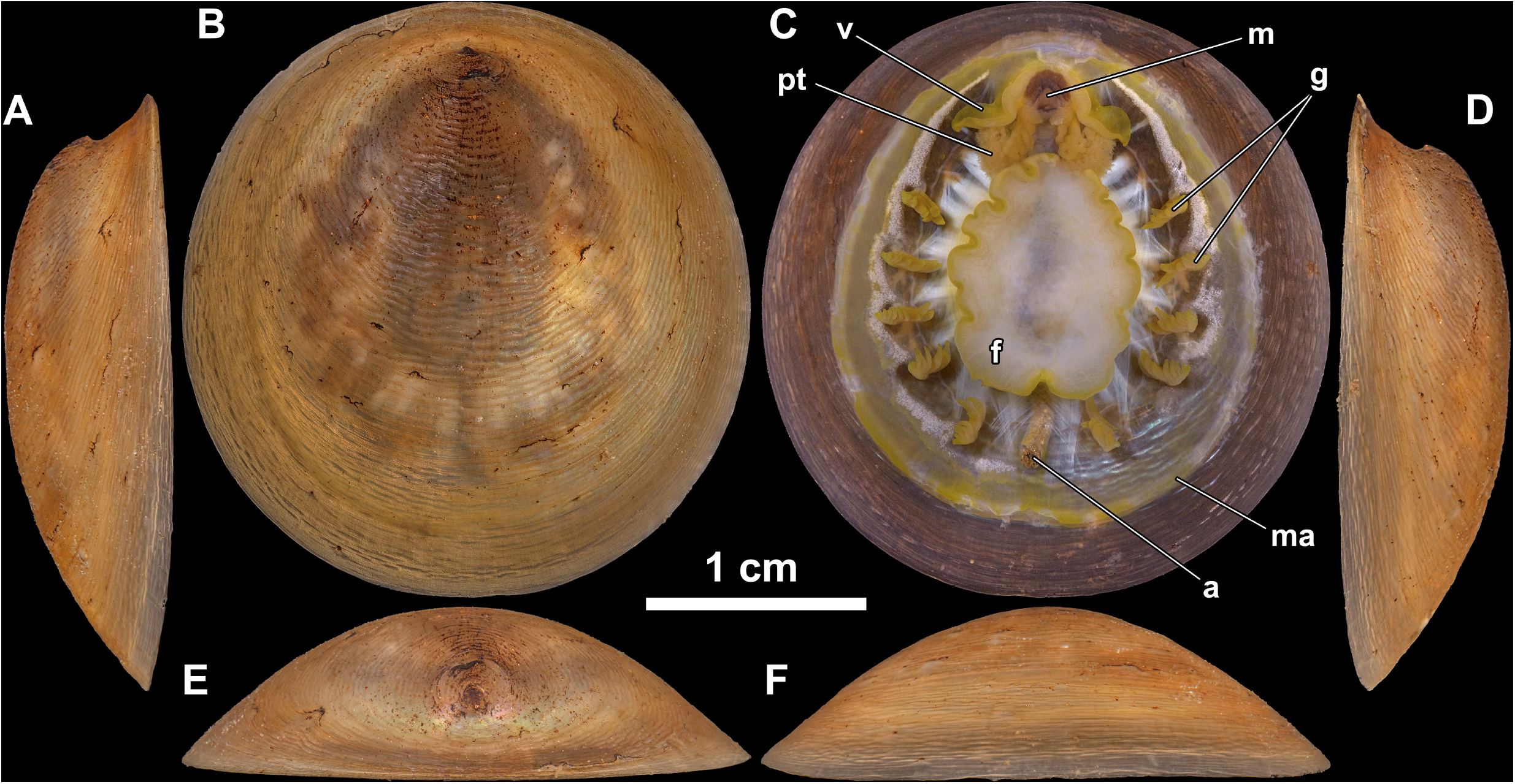
The newly collected specimen of *Neopilina galatheae* photographed from six different angles, anterior is upwards. A: view from left side; B: dorsal view; C: ventral view; D: view from right side; E: anterior view; F: posterior view. Abbreviations: a, anus; g, gills; m, mouth; ma, mantle; pt, postoral tentacles; v, velum.

**Figure 3.**
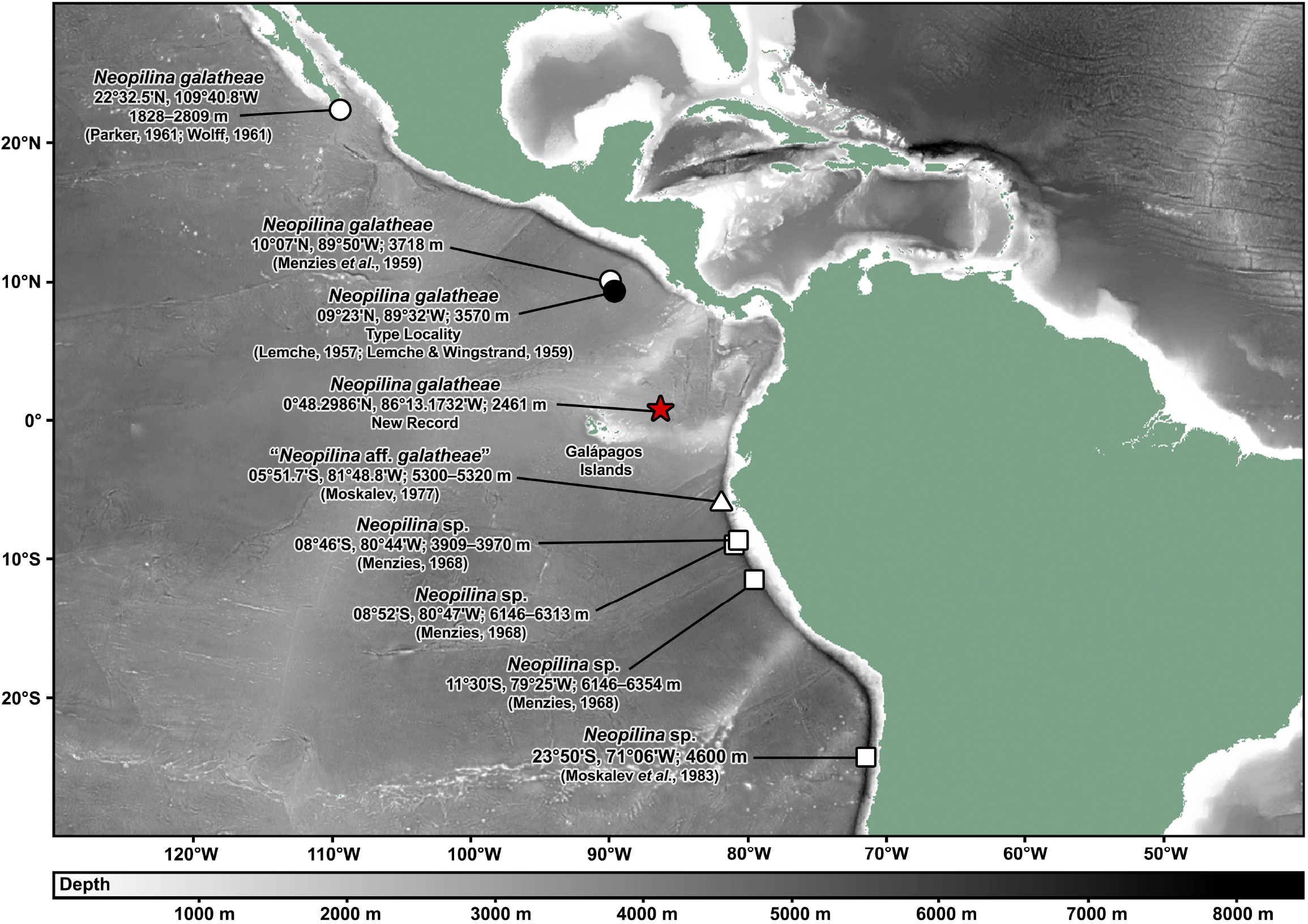
Summary map of all confirmed distribution records of *Neopilina galatheae* (closed circle for the type locality, open circle for other records, and red star for the new record herein), and eastern Pacific records of *Neopilina* with uncertain species-level identity, including “*Neopilina* aff. *galatheae*” *sensu* Moskalev, 1977 (open triangle) and *Neopilina* sp. (open squares).

The present record of the monoplacophoran *Neopilina galatheae* is the southernmost record attributable to this rare species with certainty (Lemche & Wingstrand, 1959; Menzies, 1968; Parker, 1961; Schwabe, 2008), extending its distribution approximately 1000 km south-eastwards from the type locality off Costa Rica. There are, however, several further monoplacophoran records ascribed to *Neopilina* in the eastern Pacific south of the equator. The most notable is a record from the Atacama Trench off Chile (5°51.7’S, 81°48.8’W; 5300–5320 m deep) attributed to “*Neopilina* aff. *galatheae*” (Moskalev, 1977), which is actually further south than our present new record. Unfortunately as this was only a passing mention in the footnote of a paper focusing on patellogastropod limpets with no information on size, morphology, or repository provided, this record cannot be verified. This record is geographically close to several other Atacama Trench records (5°51.7’-23°50’S, 71°06’-81°48.8’W; 3909–6354 m deep) attributed to *Neopilina* sp. (Menzies, 1968; Moskalev, Starobogatov & Filatova, 1983). Since Menzies (1968) who has previously collected *N. galatheae* (Menzies *et al*., 1959; Menzies & Layton, 1962) did not attribute these to *N. galatheae*, it is reasonable to consider these to be another, possibly undescribed, species. The Moskalev (1977) record may be an additional record of this *Neopilina* sp.; though these findings may also be a compilation of multiple species. Furthermore, Moskalev (1977) listed the *Neopilina* aff. *galatheae* as a specimen of about 20 mm length from R/V Akademik Kurchatov station K-301, but in a follow-up study led by the same author (Moskalev *et al*., 1983) the only monoplacophoran listed from this station was one *Vema ewingi* at 17.3 mm shell length. As such, it also seems likely that these two records refer to the same specimen and the ‘*Neopilina* aff. *galatheae*’ Moskalev (1977) may be a misidentified *V. ewingi*, but the slight difference in shell length between the two publications makes this inconclusive. To clarify the taxonomic affinity of these records, additional sampling at the Atacama Trench is warranted. Figure 3 presents a summary of these relevant *Neopilina* records from the eastern Pacific.

The new specimen of *N. galatheae* was found alive on a solidified basaltic lava flow, in an area covered by a recent (2005-2012) eruption event (Shank *et al*., 2012) and therefore devoid of thick sediment. This is in strong contrast to other known habitats for this species, including muddy clay surfaces (Lemche & Wingstrand, 1959), greenish ooze (Menzies *et al*., 1959), and sandy mud rich in organic matter (Parker, 1961). Nevertheless, the only other *in situ* seafloor imagery of *Neopilina* was also taken on a basalt surface on a seamount (Sigwart *et al*., 2019). Together with our observations, this suggests *Neopilina galatheae* (and *Neopilina* in general) may be able to inhabit both hard and soft substrata. Unlike patellogastropod limpets which avoids sedimented bottoms, monoplacophorans happily transverse sediment (Sigwart *et al*., 2019) which is perhaps an indication of their capacity to inhabit multiple substrate types. As trawled gears and box cores are deployed on soft bottoms to avoid damaging them, the sampling method explains the bias of the early records on soft bottoms (Parker, 1961). This, of course, is also supported by numerous findings of small-bodied monoplacophorans taken on hard substrates, particularly manganese nodules (Lowenstam, 1978; Wiklund *et al*., 2017) and the larger monoplacophoran *Adenopilina adenensis* (Tebble, 1967) whose collection substrate was not given in the description (Tebble, 1967) but the cruise report suggests a hard substrate (Laughton, 1967).

Monoplacophorans are generally regarded as detritivores, and in this specimen we found faecal material similar in colouration to the surrounding thin sediment (Figure 2), agreeing with previous studies (Menzies *et al*., 1959; Sigwart *et al*., 2019). We did not, however, see clear grazing trackways associated with or around the animal as was sighted in a previous *Neopilina* encounter (Sigwart *et al*., 2019). One estimate of *N. galatheae* population density based on trawl area indicated an exceedingly low average density of one individual per 22,000 square meters (Menzies *et al*., 1959). The finding of 10 specimens in a single trawl, however, would suggest local aggregations (Menzies *et al*., 1959); a scenario also hinted by the Samoan sighting of *Neopilina* based on the high density of feeding trackways (Sigwart *et al*., 2019). Although it is possible that this particular individual was not feeding, another possibility is that the glassy, black lava substrate made it difficult to see feeding trails. *Neopilina galatheae* has also been suggested to feed on xenophyophores (Tendal, 1985). We did see numerous xenophyophores during the ROV dive, but we did not see *N. galatheae* feeding on them.

This *in situ* rediscovery of the first monoplacophoran *Neopilina galatheae* provides new insights on its distribution and ecology, and also a valuable specimen for future studies. We now know that fossil monoplacophorans are likely a collection of distantly related lineages that happened to have a similar shell morphology (Haszprunar & Ruthensteiner, 2013), but the Paleozoic family Tryblidiidae within Monoplacophora have eight muscle scars strongly resembling living species and are likely directly related to them (Ponder *et al*., 2020). Molecular phylogeny, however, has suggested that living monoplacophorans may have diverged only in the Late Cretaceous (Kano *et al*., 2012) – leaving a gap of about 300 million years in their record. The new *N. galatheae* specimen is likely the first large-bodied living monoplacophoran specimen collected in decades (Schwabe, 2008), and potentially also the first such specimen available for genomics work (Kocot *et al*., 2020). It is hoped that future data from this specimen will shed bright light on the evolutionary history of these enigmatic “living fossils”.

## Acknowledgements

I thank the captain and crew on-board R/V Falkor (too) during the research cruise FKt231024 (‘Project Zombie: Bringing dead vents to life – Ultra Fine-Scale Seafloor Mapping’) for their great support of the scientific activities. I extend this thanks to the ROV SuBastian support team, most notably Jason Rodriguez who collected the monoplacophoran individual examined herein. All on-board scientists of the cruise FKt231024 are gratefully acknowledged, especially the chief scientist John W. Jamieson (Memorial University of Newfoundland) for his diligent execution of the research cruise. The research cruise FKt231024 on-board R/V Falkor (too) was funded by Schmidt Ocean Institute. Sigrid Hof and Sandra Müller (Senckenberg Museum, Frankfurt) were extremely helpful in procuring key literature for this study. Miwako Tsuda (JAMSTEC) assisted with lab work.

## Data Availability

All relevant data are provided within the manuscript or the Supplementary Video available on Figshare (Chen, 2024), DOI: 10.6084/m9.figshare.24503623. The *Neopilina galatheae* specimen is deposited in Senckenberg Natural History Museum, Frankfurt (SMF 373198) and its COI (mitochondrial cytochrome oxidase *c* subunit I) barcode (Folmer *et al*., 1994) has been made available on NCBI GenBank under the accession number PP352643.

